# A low-cost, open-source evolutionary bioreactor and its educational use

**DOI:** 10.1101/729434

**Authors:** Vishhvaan Gopalakrishnan, Dena Crozier, Kyle J. Card, Lacy D. Chick, Nikhil P. Krishnan, Erin McClure, Julia Pelesko, Drew F.K. Williamson, Daniel Nichol, Soumyajit Mandal, Robert A. Bonomo, Jacob G. Scott

## Abstract

The morbidostat automatically adjusts antibiotic concentration as a bacterial population evolves resistance. Although this device has advanced our understanding of the evolutionary and ecological processes that drive antibiotic resistance, no low-cost and open-source systems are available for educators. Here, we present the EVolutionary biorEactor (EVE), an accessible alternative to other morbidostats for use in low-resource classrooms that requires minimal engineering and programming experience. We first compare our system to others, emphasizing how it differs in design and cost. We then describe how we validated the EVE by evolving replicate *Escherichia coli* populations under chloramphenicol challenge and comparing our results to those in the published literature. Lastly, we detail how high school students used the EVE to learn about bacterial growth and antibiotic resistance.

## Introduction

Continuous culture vitalized microbiological research. Bioreactors automatically introduce sterile growth medium and remove unused nutrients, cells, and metabolic byproducts to maintain microbial populations over time under steady-state conditions (***Novick and Szilard, 1950***). The first bioreactor, the chemostat, was introduced in the mid-20th century to maintain populations at a particular specific growth rate by sustaining a constant rate of nutrient exchange (i.e., dilution rate) (***Novick and Szilard, 1950***). The study of populations under these environmental conditions made physiological and biochemical analyses more tractable and led to the development of quantitative models that describe microbial growth (***Hoskisson and Hobbs, 2005***).

The growing public health concern of antibiotic resistance has prompted a renewed focus on continuous-culture techniques to improve our understanding of the evolutionary and ecological dynamics underlying this phenomenon. For example, the morbidostat is a bioreactor that regulates bacterial growth rate by measuring population cell density at fixed time intervals. An antibiotic is then introduced into the medium when the population size exceeds a predetermined threshold and the growth rate is greater than zero (***Toprak et al., 2012***). The morbidostat therefore adjusts the selective pressure to maintain near-constant growth inhibition as the population evolves resistance. Although this device has advanced our understanding of the repeatability of resistance evolution in bacteria (***Toprak et al., 2012**; **Dößelmann et al., 2017**; **Yoshida et al., 2017**; **Leyn et al., 2021)*** and temperature stress response in yeast (***Wong et al., 2018***), the lack of open-source software and high commercial cost may limit its widespread use, particularly in educational settings.

Implementing this technology in a high school or college curriculum has many benefits. First, educators often use passive, lecture-based instruction to teach that evolution is a gradual process that occurs over long timescales. In contrast, in-classroom morbidostat use enables authentic, inquiry-based research experiences in which students combine principles of hypothesis generation and appropriate experimental design with observations of the real-time adaptation of microbial populations. This active learning approach improves student understanding of bacterial growth dynamics and how natural selection acts on heritable variation to promote drug resistance. Thus, the use of a morbidostat in the classroom complements existing inquiry-based laboratory curricula (***Cooper et al., 2019***). Second, device assembly teaches students critical engineering and computer programming skills, including working with modern computer architectures, operating systems, programming languages, and electronics that sense and interact with the physical environment. We developed a morbidostat, the EVolutionary biorEactor (EVE), with these goals in mind.

The EVE’s low production cost and open-source design uniquely position it to advance evolutionary understanding in the classroom. We constructed the EVE by replicating and expanding upon the methodology outlined by Toprak and colleagues (***Toprak et al., 2012**, **2013***), focusing on lowering costs and improving build transparency. Our platform supports multiple culture units (CUs) that each run independently with precise control of growth medium exchange, cell density measurement rates, and drug addition. One can construct the EVE for $115-200 using printed circuit board (PCB) and 3D-printed component files, circuit diagrams, software, and complete build instructions in our GitHub repository (***Gopalakrishnan, 2022***).

In this Feature Article, we compare the EVE to other morbidostats with particular attention to its unique characteristics. We then document the EVE’s use by high school students in Dr. Lacy Chick’s Advanced Placement (AP) Biology classroom at Hawken School in Gates Mills, Ohio. We developed a curriculum for the evolutionary unit of the AP Biology course that detailed the desired learning outcomes, assessment evidence for those outcomes, and a learning plan. The students then assembled an EVE device from provided materials and performed experiments to assess bacterial population growth over time in permissive (i.e., drug-free) and selective media. By direct observation and real-time measurement, they learned that bacterial populations could steadily evolve resistance when subjected to increasing drug concentrations. Therefore, they established that the EVE is an accessible alternative to other bioreactors for use in low-resource classrooms, and where instructors and students have limited engineering and programming experience.

## The EVolutionary biorEactor

The EVE is functionally similar to other small automated bioreactors (***Toprak et al., 2012**; **Dößelmann et al., 2017**; **Wong et al., 2018***). In summary, several bacterial cultures are grown simultaneously in small glass vials under well-mixed, drug-selective conditions (Figure 1A). A control algorithm maintains these conditions by(i) supplying power to fans (with attached magnets) that rotate magnetic stir bars; (ii) introducing fresh medium into each culture at a constant rate through the activation of peristaltic pumps; (iii) monitoring population growth via absorbance measurements taken from paired light-emitting and photo-sensitive diodes; and (iv) introducing selective medium when these absorbance measurements exceed a user-defined value. Nonetheless, the EVE differs from other bioreactors in its hardware, software, and production costs. We detail these aspects below.

**Figure 1.**
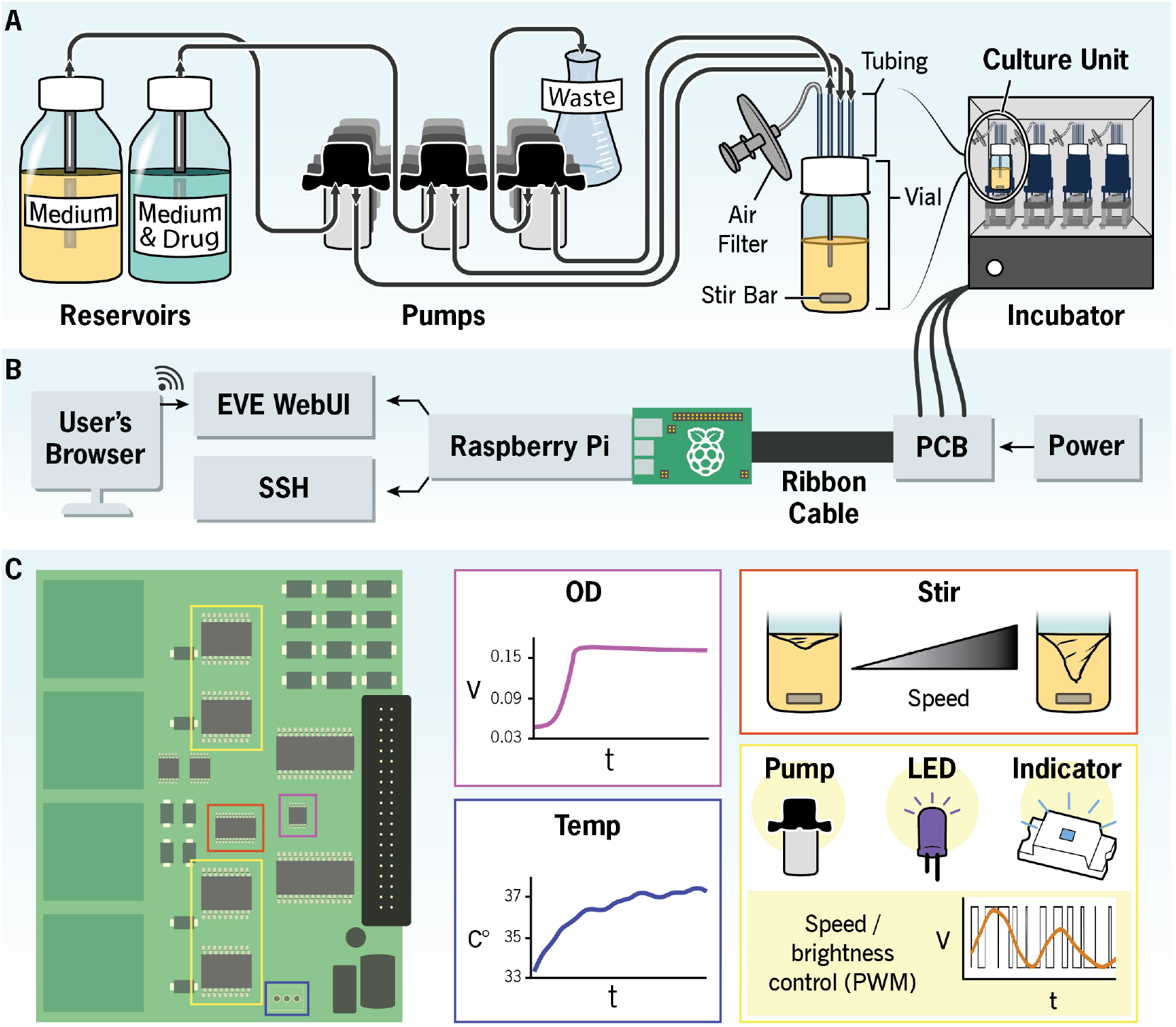
Schematic illustration of the EVE hardware and software architecture. **(A)** Reservoirs containing permissive (i.e., drug-free) and selective growth media are each connected to a culture unit (CU) through silicone tubing. Peristaltic pumps control the rate at which the media are added to the evolving bacterial population, and an additional pump removes waste. Multiple CUs can fit into an incubator, allowing users to simultaneously evolve several independent replicate populations. **(B)** A Raspberry Pi interfaces with a custom printed circuit board (PCB) to monitor culture growth and coordinate the addition and removal of growth media and waste. The user controls the EVE hardware with a web application. **(C)** The PCB chips measure voltage and temperature. These voltage measurements are proportional to optical density and thus are a way to estimate population size. The chips also control the fluidic pumps, diodes, onboard indicators, and stirring speed The PCB diagram illustrates where these chips are located on the board.

### Hardware and software

In contrast to other devices that use an external system to process data (***Toprak et al., 2012**; **Wong et al., 2018)***, the EVE uses a primary onboard Raspberry Pi to execute the software and serve as a bridge between the user interface and a PCB (Figure 1 B, C). This design keeps costs lowwhile allowing the device to be self-contained in environments without internet access. One can also construct the EVE with a breadboard. Breadboards give users more flexibility to modify the hardware but come at the cost of increased assembly time, required prior experience, and device footprint. In either case, circuit diagrams, 3D-printed component files, complete build instructions, and a part list are in our GitHub repository (***Gopalakrishnan, 2022***).

One may access population growth and temperature measurements, edit configuration files, define experimental parameters, and monitor and control experiments remotely with our free, open-source software run inside a web browser. Users install this software by downloading the pre-compiled Raspberry Pi image orexecutingthe installation script directly from our GitHub repository (***Gopalakrishnan, 2022***). Data can be saved to network locations mounted to the Pi’s file system or an attached USB device.

We wrote the software in Python, a common programming language with a broad user base and community support. One does not need to be familiar with this language to install the software, edit experimental parameters, or run the pre-configured experimental algorithm. However, working knowledge of Python is necessary to create custom drug selection algorithms. In contrast, other morbidostat systems use proprietary (***Wong et al., 2018***) or third-party (***Toprak et al., 2012***) software that might be inaccessible or cost-prohibitive for educators.

### Production cost

We designed the EVE to be cost-effective. The total cost varies between $115 to $200 to build a system that performs simple growth or evolution experiments in triplicate with a negative control. These estimates include all parts except the incubator, glassware, and 3D printer. The circuit board design can be downloaded from our repository and sent to a PCB manufacturer for printing and assembly; in our case, we purchased fully-assembled PCBs for approximately $42 per board, including the power supply. Users may further decrease costs and increase accessibility in low-resource classrooms by incubating bacterial cultures at room temperature and using a pressure cooker to sterilize growth media instead of an autoclave.

## Validation

We validated the EVE in two ways. First, we examined voltage measurement variability across seven independent CUs and compared these results with previously reported measurements ***(Toprak et al., 2012)***. The average variability was 8.0% (mean ± sd = 2.93 ± 0.23), comparable to the 7.5% of Toprak and colleagues. Second, we experimentally evolved bacterial populations and again compared the results to those of Toprak et al. (2012). We revived *E. coli* ATCC 25922 from a frozen stock by overnight growth in Mueller-Hinton broth (MHB) (Sigma). We inoculated 100 μL of this culture into three separate CUs containing 12 mL of MHB. We then prepared the selective growth medium by diluting a chloramphenicol stock solution to 40 μg/mL in MHB. The selective and permissive media were connected to the CUs, which were incubated at 37°C with constant stirring (225 rpm). Samples were taken from each culture’s effluent waste every 12 h and frozen at –80°C with 15% glycerol as a cryoprotectant.

We estimated each sample’s half-maximal inhibitory concentration (IC_50_) following the broth microdilution method (***Wiegand et al., 2008***) and by fitting a Hill function to the resulting optical density data (***Maltas and Wood, 2019***). The three ancestral populations began the experiment with an IC_50_ of 4 μg/mL, and chloramphenicol resistance increased to 9.3 - 13.9 μg/mL. These results are comparable to Toprak et al. (2012), where resistance increased, on average, to approximately 15 μg/mL over the same time period (Figure 2).

**Figure 2.**
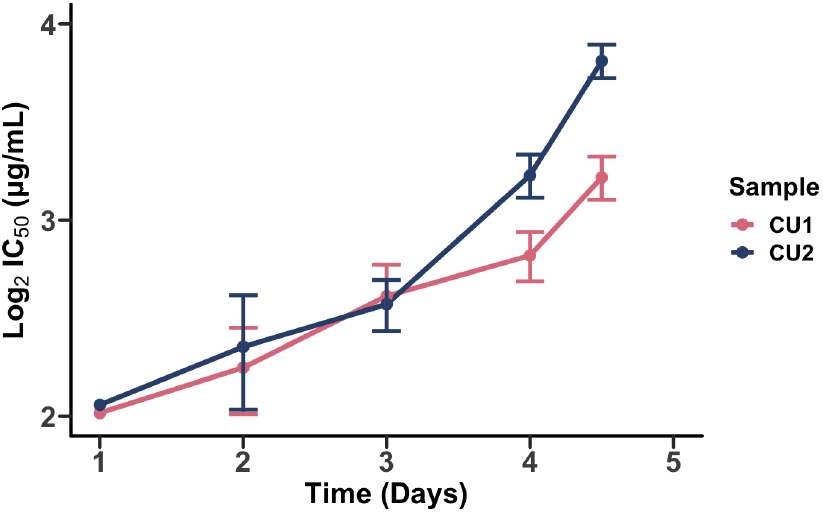
The evolution of chloramphenicol resistance overtime. Half-maximal inhibitory concentration (IC_50_) values are shown on a log_2_-transformed scale to reflect that antibiotic concentrations were tested across several two-fold dilutions. Points represent the estimated IC_50_ determined by fitting a Hill function to OD_600_ measurements across this dilution series after 18 h of population growth. Error bars represent the variance in IC_50_.

## Educational Use

The EVE is well-suited for classroom settings because it was designed in collaboration with educators and tested by high school students. Here, we discuss our curriculum design, the students’ experience using the EVE, and its potential applications in other educational contexts.

We first examined the AP Biology curriculum and developed a unit plan to complement this program. The AP curriculum introduces several key learning objectives about the importance of phenotypic variation, how natural selection acts on this variation, and how this phenomenon affects populations over time. The AP instructional model also emphasizes that students use supporting resources and perform appropriate experiments to build and strengthen their conceptual understanding of these objectives. More generally, for most biology classrooms, bioreactor experiments introduce basic microbiology techniques, biotechnology, and data analysis to students. We therefore developed our unit to facilitate student understanding of population growth dynamics and trait evolution through independent experimentation with the EVE. To reinforce this objective, we asked the students to consider how bacterial populations respond to changing environments, such as introducing antimicrobial agents.

Two high school students followed the instructions in our GitHub repository ***(Gopalakrishnan, 2022)*** to assemble the EVE bioreactor with 3D-printed equipment, a fabricated circuit board, and a Raspberry Pi microcomputer with the necessary pre-installed software. Then three AP Biology students, working as a team, calibrated the device and evolved a population of *E. coli* ATCC 25922 to increasing chloramphenicol concentrations over several days (Figure 3). The population initially expanded in size, decreased after the introduction of the antibiotic into the culture unit, and later rebounded as drug-resistant variants rose in frequency (Figure 3H). From these data, the students calculated growth rates and predicted how these rates would change with varying drug selective pressures. This authentic research experience introduced the students to standard microbiology practices and concepts, including how to prepare drug solutions and use sterile technique to maintain bacterial cultures, when logistic growth models apply, and the relationship between light absorbance and cell number (i.e., Beer’s law). We share their testimonials in Box 1.

**Figure 3.**
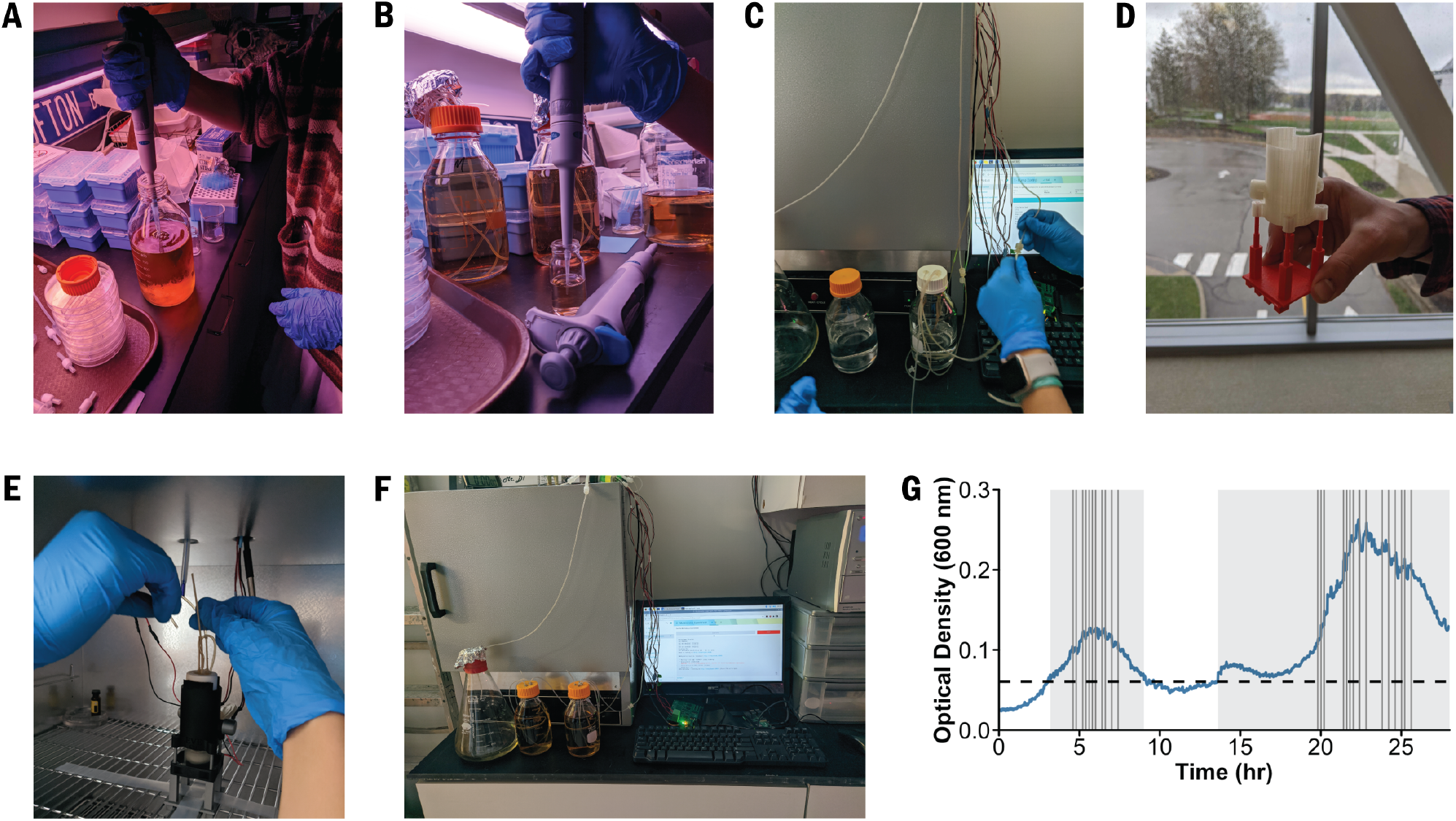
High school students built the EVE and performed an evolution experiment. **(A)** They prepared the selective growth medium and **(B)** inoculated *E. coli* into a culture unit (CU) containing the permissive medium. **(C)** The system was sterilized with bleach and ethanol solutions. **(D)** Before the experiment, the students printed a CU stand in the Hawken School’s MakerSpace. **(E)** The media and waste reservoirs were attached to the CU by silicon tubing, and then **(F)** the students began the experiment. **(G)** Bacterial growth and inhibition in the EVE over 28 h. The grey boxes and vertical lines indicate when the EVE control algorithm added the permissive and selective media into the culture units, respectively.

### Box 1. Student Experiences

The bioreactor experiment was a very valuable learning experience. Learning the evolution of bacterial growth against drug resistance deepened my understanding of the wonders of the evolution unit in my current biology class. Through our hands-on experience, I have learned many new terms and biotechnology techniques including pipetting, working with bacterial cultures, and learning the Beer’s Law. This was a complex but also at the same time simple lab to do, and I think it would definitely be beneficial for students to learn more about bacterial evolution and practice important skills such as calculating concentrations and analyzing data.

*— Lillian Fu*

The Bioreactor experiment was very informational for me; I learned how bacterial growth works and how drug resistance plays a role in it through my first hand experience with being able to actually work with the bacterial cultures, drugs, pipettes, and EVE. I think this was a fun project to do, and it would definitely be an easy and helpful project for other high school students to do while learning about bacterial growth and evolution. Students can also utilize and strengthen skills such as analyzing data and calculating different concentrations. Overall, I think this project was accessible and simple but still very interesting and informative.

*— Grace Shum*

Notably, the students adjusted the experimental protocol to account for classroom limitations. For example, the Hawken School does not have Bunsen burners or standing incubators. The students therefore created a spartan workspace and disinfected all surfaces with 70% ethanol, visually inspected the growth medium to ensure that it remained free of contamination, and incubated bacterial cultures in an oven set to approximately 30 - 40°C. Moreover, although this particular high school has autoclaves, there are several alternative ways that users can sterilize glassware and media, including using liquid chemicals and microwaves. The Centers for Disease Control and Prevention describe some of these methods in greater detail (***CDC, 2016***).

One could also use the EVE in other educational contexts (i.e., college laboratory classrooms) or outside the classroom. Since the EVE functions through a combination of engineering and biology, project work within clubs may offer unique experiences to students wishing to learn about how multiple disciplines interact. These projects could combine construction and experimentation for a seamless educational experience. Interested parties may acquire non-pathogenic *E. coli* K12, growth medium, and antibiotic powders through the Carolina Biological Supply Company or a similar vendor.

The EVE has also been implemented in several other contexts. For instance, the EVE is being used to study bacteriophages in continuous culture at the University of Exeter in England, and used to develop research bioreactors in a French biotech company. Additionally, undergraduate students used its design to build their own custom morbidostat as part of the International Genetically Engineered Machine (iGEM) competition (***GEM, 2019***). Their manual represents an example of what the EVE would look like in a college setting.

## Future Directions

In addition to its educational utility, the EVE is an ideal system to address questions of evolutionary repeatability. For instance, one might examine whether correlated drug responses are conserved across time as populations evolve under single- or multi-drug selection (***Nichol et al., 2019***). Although the current EVE system can only introduce one drug solution into the growth medium, we are currently designing a PCB that will allow the simultaneous or sequential addition of multiple drugs. Second, users could substitute the existing hardware with LEDs and photodiodes corresponding to fluorescence proteins’ excitation and emission frequencies. This hardware alteration would allow bacterial head-to-head competitions without periodic sampling and cell enumeration.

## Conclusion

In this Feature Article, we introduced the EVE, a bioreactor that is uniquely suited for use in educational settings because of its economical, accessible, and open-source design. High school students demonstrated these aspects by building the EVE and performing a simple evolution experiment in the classroom. In summary, we are optimistic about a future where more classrooms use the EVE and other platforms to spur student interest in experimental biology and technology.

