## supplemental documents for "A low-cost, open-source evolutionary bioreactor and its educational use"

### Supplementals

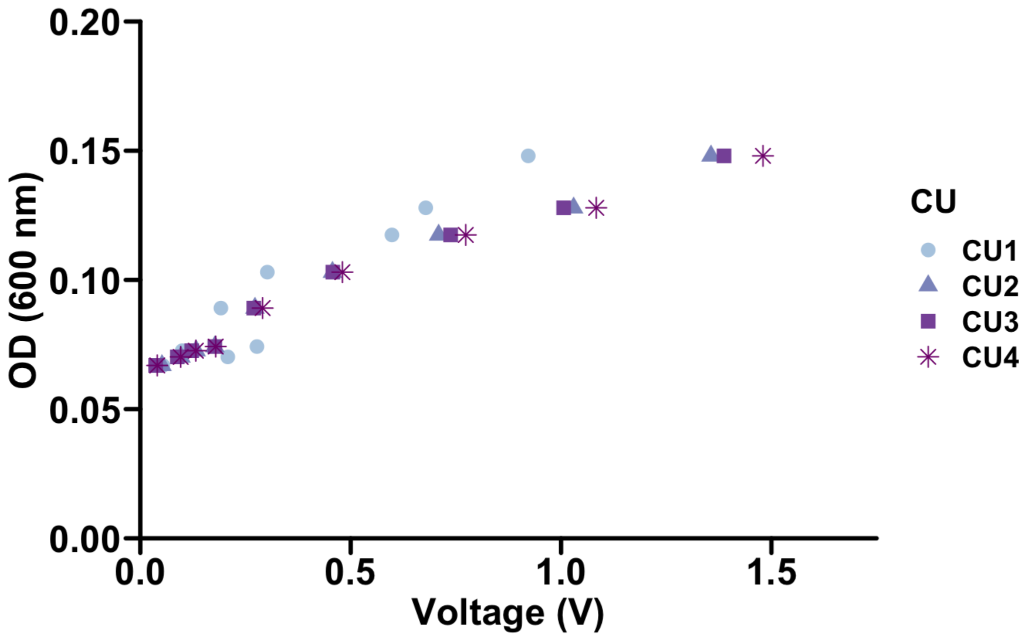


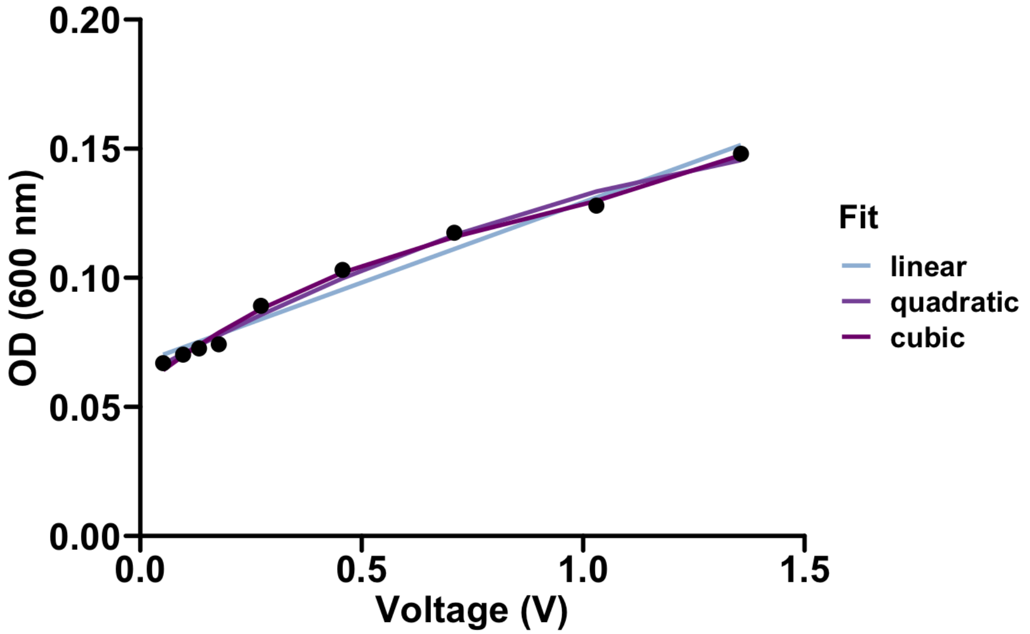


**Figure 1.** Calibration of culture optical density (OD) with reported voltage. (Top) Culture units behave similarly when measuring voltage. (Bottom) Various regression methods were used to quantify the relationship between the OD and voltage. $R^2$ values of 0.97, 0.99, and 0.99 were calculated for the linear, quadratic, and cubic fits respectively. We used a linear model to translate voltage to optical density in our experiments.

**CURRICULUM UNIT PLAN**

*The Department of Translational Hematology and Oncology Research at the Cleveland Clinic*

| | | |
|---|---|---|
| **Course: Biology/AP Biology** | | **Grade Level: 11-12** |
| **Unit Title: Biotechnology and Evolution** | | **Length of Unit: 3 Weeks** |
| **Concept: Grow harmless bacteria in a self-built automated culture device which monitors growth in real time. Optionally introduce bactericidal agents (i.e., watered down toothpaste) and watch bacteria die and recover from the insult.**<br><br>**Unit Summary: Students will learn the concepts involved with evolution via natural selection and trait variation within a population. Through this, they will be able to manipulate aspects of bacterial growth and decline through inquiry-based labs and population growth models. These multi-week lessons may be used to cover topics in biotechnology, evolution, antibacterial resistance, population growth, or the scientific method and variable manipulation.** | | |
| **Stage 1- Desired Results** | | |
| | **Transfer** | |
| | *Students will be able to independently use their learning to experiment with population growth models and trait selection in response to environmental stimuli.* | |
| | **Meaning** | |
| | UNDERSTANDINGS<br>*Students will understand that…*<br><br>By manipulating environmental conditions, population growth will change to reflect | ESSENTIAL QUESTIONS<br>*Students will be able to answer...*<br><br>What happens to the bacterial populations when conditions (i.e., variables) are altered? |

| | | |
|---|---|---|
| | optimal conditions.<br><br>Optimal conditions vary for different types of bacteria. Population growth can be calculated and modeled based on predictions of the environment and knowledge of optimal growth rates.<br><br>Bacteria can change through generations via natural selection in response to selection pressures imposed by the environment.<br><br>Distributions of traits change and can be selected for over multiple generations within a population. | What are the *optimal* environments which would allow for different types of bacterial populations to grow?<br><br>What are *suitable* environments which would allow for different types of bacterial populations to grow?<br><br>What is the growth rate of bacterial populations under different conditions?<br>How do different concentrations of bactericidal agents affect traits and bacteria populations?<br><br>Do bacterial populations follow density-dependent or -independent growth rates? |
| | **Acquisition** | |
| | *Students will know…*<br><br>The process of evolution by natural selection.<br><br>How to calculate population growth/decline. | *Students will be skilled at…*<br><br>Manipulating variables to determine optimal bacterial growth.<br><br>Calculating growth rates and declines, and predicting outcomes based on differing selection pressures.<br><br>Adjusting the environmental conditions to get different desired population outcomes. |

| Stage 2- Evidence | |
|---|---|
| **Evaluation Criteria** | **Assessment Evidence** |
| Task Rubric | PERFORMANCE TASK(S):<br>Assemble a Bioreactor from provided materials, or (if applicable) print and assemble materials from publicly-available software (i.e., github). Design and implement an experiment with independent and manipulated variables. Monitor and predict bacterial evolution by manipulating selection pressures to see how populations recover and vary from changing environments. |
| | COMMON UNIT ASSESSMENT<br>Unit projects with presentations which will include comparative graphs and a literature review of their bacteria. |
| | OTHER EVIDENCE<br>Track, graph, and analyze bacterial growth and decline through time over multiple labs. Determine how manipulation of variables can lead to differences in population growth rates. |

| Stage 3- Learning Plan |
|---|
| *Summary of Key Learning Events and Instruction*<br><br>Students get hands-on experience with experimental design and biotechnology, learn to manipulate variables, monitor and predict outcomes, and see evolution in action over a 3-week lesson spanning multiple unit concepts in both the General and AP Biology curricula. |
